# Droplet and fibril formation of the functional amyloid Orb2

**DOI:** 10.1101/2021.02.19.430336

**Authors:** Kidist Ashami, Alexander S. Falk, Connor Hurd, Samridhi Garg, Silvia A. Cervantes, Anoop Rawat, Ansgar B Siemer

## Abstract

The functional amyloid Orb2 belongs to the cytoplasmic polyadenylation element binding (CPEB) protein family and plays an important role in long-term memory formation in *Drosophila*. The Orb2 domain structure combines RNA recognition motifs with low complexity sequences similar to many RNA binding proteins shown to form protein droplets via liquid-liquid phase separation (LLPS) in vivo and in vitro. This similarity suggests that Orb2 might also undergo LLPS. However, cellular Orb2 puncta have very little internal protein mobility and Orb2 forms fibrils in *Drosophila* brains that are functionally active indicating that LLPS might not play a role for Orb2. In the present work, we reconcile these two views on Orb2 droplet formation. We show that soluble Orb2 can indeed phase separate into protein droplets. However, these droplets have either no or only an extremely short-lived liquid phase and appear maturated right after formation. For Orb2 fragments that lack the C-terminal RNA binding domain (RBD), droplet formation is a prerequisite for fibril formation of an otherwise stable monomeric Orb2 solution. Solid-state NMR shows that these fibrils have additional well ordered static domains beside the Gln/His-rich fibril core. Further, we find that full-length Orb2B, which is by far the major component of Orb2 fibrils in vivo, does not transition into cross-β fibrils but remains in the droplet phase. Together, our data suggest that phase separation might play a role in initiating the formation of functional Orb2 fibrils.

## 1. Introduction

Orb2 is a cytoplasmic polyadenylation element binding (CPEB) protein that can form functional cross-β (amyloid) fibrils with a regulatory role for long-term memory (LTM) formation in *Drosophila* [1]. In its monomeric form, it promotes the deadenlyation of target messenger mRNA. When aggregating into cross-β fibrils, it becomes an activator of the polyadenlyation and thereby the translation of mRNAs resulting in a stabilization of memories past 48 h [2, 3, 4, 5]. Orb2 has two isoforms, Orb2A and Orb2B, which both share two C-terminal RNA recognition motifs (RRMs), a C-terminal zinc finger, a central Gly-rich region, and a Gln/His-rich domain that forms the core of Orb2 fibrils (see Figure 1) [2, 6].

**Figure 1:**
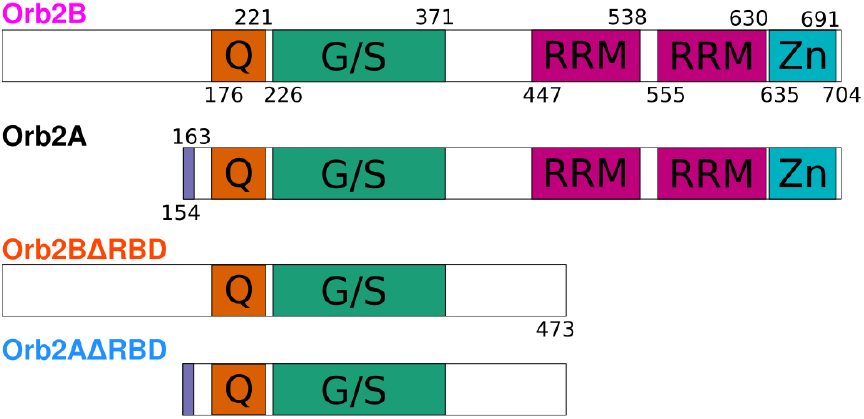
Domain structure of Orb2 isoforms and fragments used in this study. The glutamine-rich region (Q, orange), glycine/serine-rich region (G/S, green), RRM domains (magenta), and zinc-finger domain (cyan) common to both Orb2A and Orb2B are highlighted as well as the unique first 9 amino acids of Orb2A (purple), which are essential for LTM.

In vivo, Orb2A is of relatively low abundance and only increases in concentration upon synaptic stimulation [7]. However, its presence is essential for the aggregation of the common isoform Orb2B [3, 4]. Orb2A has 9 unique N-terminal residues that can form cross-β fibrils on their own [8] and whose deletion or mutations prevents Orb2 aggregation and LTM formation [4]. The deletion of Orb2A’s C-terminus has no phenotype [3, 9]. The N-terminus of Orb2B is Ser/Gly-rich and of unknown function. Hervás and co-workers recently determined the structure of the Orb2 fibril core, which located in the Gln/His-domain, using cryo-EM [6].

The domain structure of Orb2 that combines low complexity sequences with RNA binding domains is reminiscent of a whole class of RNA binding proteins, such as FUS, that are able to undergo liquid-liquid phase separation (LLPS) [10, 11, 12]. LLPS is often of functional relevance e.g. for RNA processing in the case of (stress) granules [13], or for neurotransmitter release and postsynaptic signal transmission in neurons [14, 15]. Additionally, other Gln-rich proteins such as huntingtin exon-1 (HTT_ex1_) or Whi3 have been reported to undergo LLPS [16, 17]. When observed via light microcopy, LLPS manifests as droplet or puncta formation. These droplets, while relatively fluid at first, can mature over time which results in the decreased ability to fuse with other droplets and low fluorescence recovery after photobleaching (FRAP) [18, 19]. Structurally, this maturation is caused by reduced protein diffusion in the droplet phase, which becomes a rigid protein glass. These mature droplets can in cases such as FUS mutants or HTT_ex1_ be the nucleation point of cross-β fibril formation [20, 16]. Taken together, these findings suggest that Orb2 could also undergo phase separation, which could have a potential role for its function, or its aggregation into cross-β fibrils, or both. In the following, we answer the question of whether or not Orb2 can undergo phase separation by demonstrating that it indeed can. Further, we characterize droplet maturation and fibril formation of different Orb2 fragments to understand the relationship between Orb2 droplet and fibril formation.

## 2. Results

To determine if Orb2 can phase separate in vitro, we studied full length Orb2B, and Orb2A and Orb2B fragments without the C-terminal RNA binding domain (RBD), which we refer to as ΔRBD constructs (Figure 1). We did not study full-length Orb2A because it does not require its RBD for proper function in vivo [3] and because we were not able to make it in large quantities. To test if Orb2 can undergo phase separation, we started in conditions that favored a stable monomeric state, namely 100 mM KCl, 1 M urea, 10 mM HEPES, 0.05% v/v β-mercaptoethanol, pH 7.6, here referred to as HEPES-salt-urea buffer (HSU Buffer). When left in HSU-buffer, Orb2 monomers did phase separate or form aggregates.

To introduce droplet formation, we exchanged each Orb2 construct from HSU buffer into H-Buffer (10 mM HEPES, 0.03% v/v NaN_3_ and 0.05% v/v β-mercaptoethanol, pH 6.5). Immediately upon exchange, the solutions became visibly turbid (Figure 2A). To further confirm that Orb2B, Orb2BΔRBD, and Orb2AΔRBD underwent droplet formation, we used differential interference contrast (DIC) and fluorescence microscopy of Oregon Green 488 labeled protein (Figure 2B). Round protein droplets were visible for all constructs immediately after buffer exchange. The average diameters of Orb2BΔRBD, Orb2AΔRBD, and Orb2B droplets were 1.3±1.0 μm, 2.2±1.1 μm, and 1.8±1.4 μm, respectively (see Figure S1). Morphological analysis [21] showed that the droplets had a high degree of roundness of 0.7-0.9. Besides H-buffer (i.e. low ionic strength), we found that both 10% PEG 8000 and polyA at a 1:10 RNA:protein m/m ratio were able to induce Orb2 droplet formation (see Figure S2) when added to HSU-buffer. Again, Orb2BΔRBD droplets are significantly smaller under these conditions.

**Figure 2:**
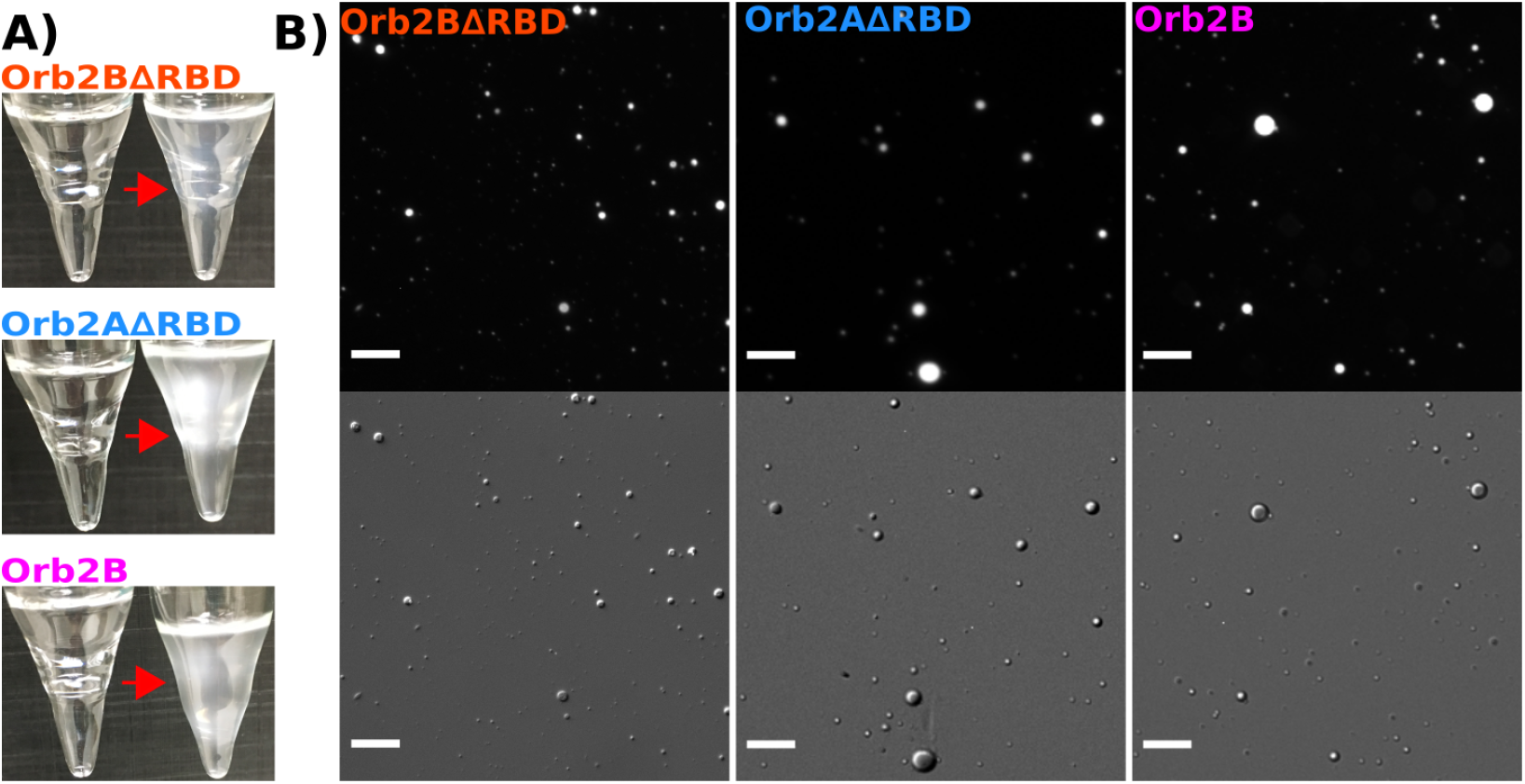
Orb2BΔRBD, Orb2AΔRBD and Orb2B can undergo liquid-liquid phase separation at low ionic strength. (A) Test tubes of Orb2 fragments before and after exchange from HSU into H-buffer. (B) Fluorescence and DIC microscopy images of droplets formed by Orb2BΔRBD, Orb2AΔRBD and Orb2B immediately after desalting from HSU-Buffer to H-buffer. Scale bars represent 20 μm.

Many protein droplets mature over time, undergoing a transition from a state in which proteins freely diffuse inside the droplet to a state with little to no protein diffusion (glass transition) [22]. To characterize protein diffusion inside droplets, we measured fluorescence recovery after photobleaching (FRAP). We bleached the center of Oregon Green 488 labeled Orb2 droplets using an argon laser and monitored the recovery by confocal microscopy for 5 minutes afterwards. To our surprise, none of the Orb2 constructs showed any degree of recovery, even though FRAP experiments were performed within 1 minute of droplet formation. These results indicate that Orb2 is static within droplets almost immediately after undergoing phase separation (Figure 3). To further confirm these results, we did 4 consecutive 10 minute FRAP experiments on the same cluster of Orb2B droplets. The first had still not recovered after the last experiment ended (Figure S3).

**Figure 3:**
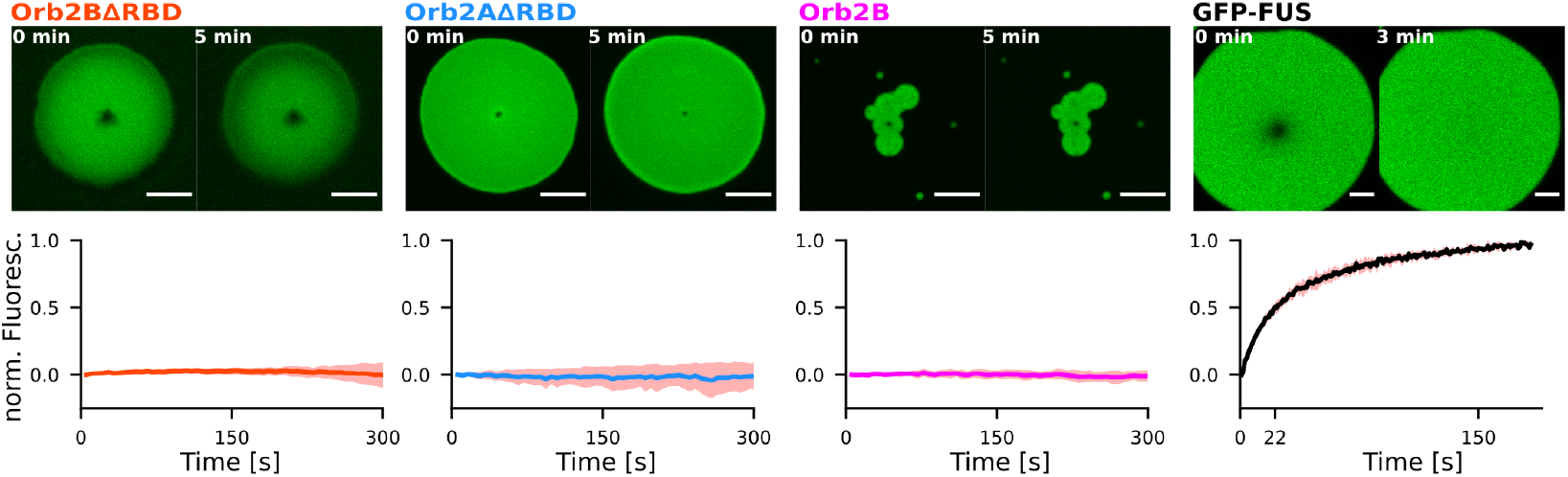
Droplets formed by Orb2BΔRBD, Orb2AΔRBD, and Orb2B show no protein diffusion via FRAP. Top: Fluorescence microscopy images right after and 5 min after bleaching. Scale bars represent 5 μm. Bottom: Average and standard deviation of the fluorescence intensity after bleaching (where 0 is the intensity after bleaching and 1 is the intensity of an unbleached region). None of the fragments showed measurable fluorescence recovery immediately after exchange into H-buffer. GFP labeled FUS, used as control, fully recovered after 3 min with 50% recovery after 22 s.

In addition, we found several clusters of Orb2B droplets with overlap between individual droplet boundaries (Figure 3 and Figure S3). For many other proteins this stage typically marks the beginning of a droplet fusion event. None of the droplets with overlapping boundaries completed the fusion process. These results support that Orb2B becomes static within droplets relatively quickly after phase separation. As a positive control, we used a N-terminally 6xHis and GFP tagged FUS construct (His-GFP-FUS) for which we measured about 50% recovery in fluorescence after 22 seconds, similar to FRAP experiments on FUS droplet reported previously [20].

For some proteins, droplet maturation is followed by cross-β fibril formation [18]. To test how Orb2 droplets evolve and if they eventually result in cross-β fibrils, we monitored Orb2 droplets over time using fluorescence microscopy, electron microscopy (EM), and Thioflavin T (ThT) fluorescence. Fluorescence microscopy and EM images of Orb2BΔRBD, Orb2AΔRBD, and Orb2B in H-buffer are shown in Figure 4. In the case of Orb2BΔRBD, the initial droplets formed halos at 12 h in our fluorescence images, which coincided with unbundled fibrils radiating out from a dark center as seen by EM. Similarly, Orb2AΔRBD droplets turned into small fibrils and droplets at 12 h. At 48 h, we observed large objects in fluorescence microscopy, which coincided with thick, bundled fibrils in our EM images. In contrast, Orb2B droplets never progressed to form fibrils, but formed larger objects that were composed of merged droplets similar to those shown in Figure 3.

**Figure 4:**
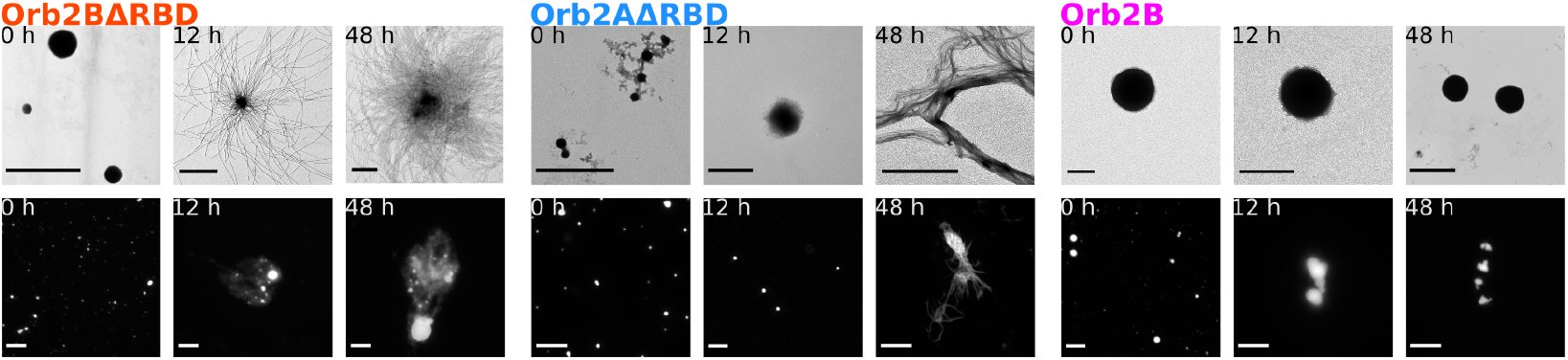
Orb2ΔRBD droplets mature into fibrils whereas Orb2B droplets do not. EM (upper row) and fluorescence microscopy (lower row) images of Orb2 constructs at 0, 12, and 48 h after exchange into H-buffer. Scale bars of EM and fluorescence microscopy images are 2 μm and 20 μm, respectively. Orb2BΔRBD and Orb2AΔRBD droplets disappear giving way to fibrils seen by EM that correlate with large fuzzy objects seen by fluorescence microscopy. In contrast, no fibrils were observed for Orb2, whose droplets seemed to merge but not fuse over time.

We attributed the disappearance of Orb2BΔRBD and Orb2AΔRBD droplets over several hours to the maturation of these droplets into cross-β fibrils (Figure 4). When examined using EM, Orb2BΔRBD fibrils were mostly unbundled, occasionally radiating out from a common center, presumably a droplet. In contrast, Orb2AΔRBD fibrils were highly bundled. As can be seen from Figure S4 both fibril types did not show any fine structure or twist at higher magnification and had an average diameter of about 18 nm.

As an additional measure of droplet to fibril transition, we measured ThT fluorescence and OD_600_ kinetics for each Orb2 construct in both HSU and H-buffer (Figure 5). We did not observe any increase in ThT fluorescence in HSU-buffer for any of the constructs. In H-buffer, Orb2BΔRBD and Orb2AΔRBD showed an increase in fluorescence in a sigmoidal shaped curve typical for cross-β fibril formation. In contrast, Orb2B showed only a negligible increase in fluoresce in H-buffer compared to HSU-buffer possibly due to light scattering from droplets. The OD_600_ curves, of Orb2BΔRBD and Orb2AΔRBD decreased in the first 20 h and increased for Orb2AΔRBD after about 25-40 h. Only a slight increase for Orb2BΔRBD was observed after about 60 h. This initial decrease is compatible with the disappearance of Orb2BΔRBD and Orb2AΔRBD droplets, whereas the later increase in scattering for Orb2AΔRBD correlates with the appearance of large, highly bundled fibrils. The OD of Orb2B stayed constant during our measurements consistent with the formation of relatively stable droplets.

**Figure 5:**
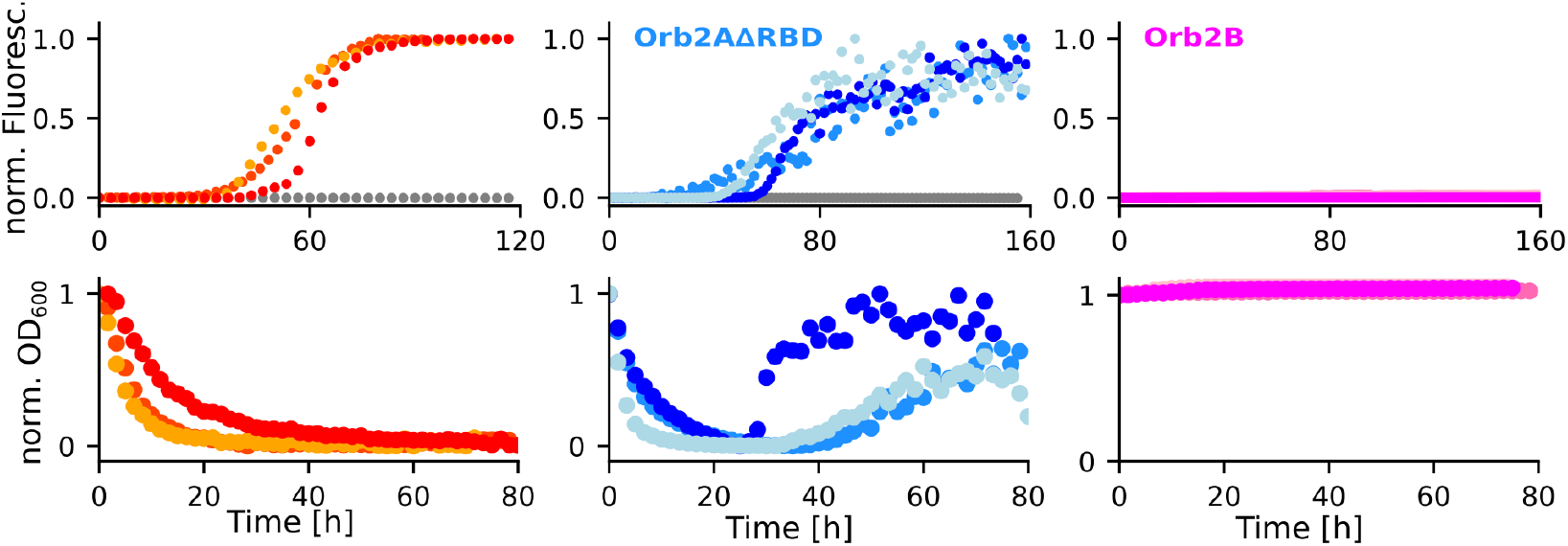
Tht fluorescence and OD measurements confirm that Orb2BΔRBD and Orb2AΔRBD droplets disappear over time giving way to cross-β fibrils whereas Orb2B droplets are stable. Top: Tht fluorescence kinetics for Orb2BΔRBD and Orb2AΔRBD were normalized to their maxima, Orb2B kinetics according to the maximum of Orb2AΔRBD because the fluorescence intensity did not increase significantly relative to background. Bottom: OD_600_ kinetics as a measure of sample turbidity. Data were normalized to their first point. Where Orb2BΔRBD and Orb2AΔRBD show an initial decrease in turbidity, Orb2 turbidity was constant compatible with stable droplet formation.

Phase separation of Orb2BΔRBD and Orb2AΔRBD ultimately led to the formation of cross-β fibrils. Next we asked, how does the structure of these fibrils compare to the cryo-EM structure of Orb2 fibrils purified from *Drosophila* brain [6] and the structure of recombinant Orb2A_1*−*88_ fibrils we investigated previously [8]? To answer this question, we expressed and purified U-^13^C-^15^N labeled Orb2BΔRBD and Orb2AΔRBD. We then prepared fibrils via phase separation for solid-state NMR measurements. One-dimensional ^13^C spectra of these fibrils are shown in Figure 6. Cross-polarization (CP) spectra detect static protein domains that in the case of cross-β fibrils often coincide with the fibril core. Refocused INEPT spectra, in contrast, are sensitive to dynamic protein regions that are often framing the static fibril core. As seen from the spectra, both Orb2BΔRBD and Orb2AΔRBD fibrils have static and dynamic domains. The aliphatic region of the CP spectra of these two fibril types overlaps relatively well indicating that both fibrils have similar structure. Both CP spectra also show His side chain resonances compatible with a His-rich fibril core. However, the small differences between the spectra indicate that their fibril core structures are not exactly the same. The more intense INEPT spectrum of Orb2BΔRBD indicates that these fibrils have larger dynamic domains.

**Figure 6:**
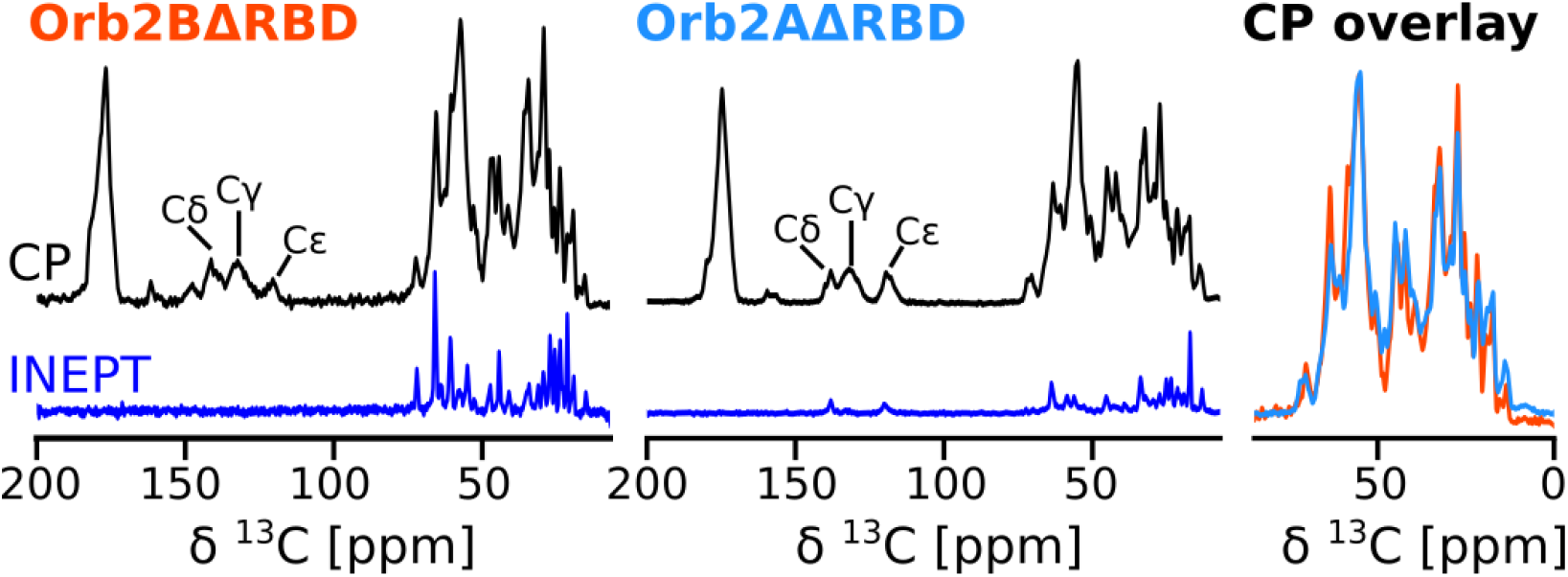
1D ^13^C spectra of Orb2BΔRBD and Orb2AΔRBD fibrils indicate that both have similar structure with larger intrinsically disordered regions found in Orb2BΔRBD. Cross polarization (CP, black) spectra detect static regions of the sample typically located in the fibril core. Histidine side chain resonances are highlighted in the aromatic region of the CP spectrum between 150 and 100 ppm. The refocused INEPT spectrum (INEPT, blue) detects highly dynamic, intrinsically disordered regions that are part of the fibril. Overlay of the two CP spectra indicates that both fibrils have similar structures.

To learn more about the amino acid residues found in the fibril core, we recorded CP-based, 2D ^13^C-^13^C DREAM (Figure 7) spectra that highlight the most immobile residues in the fibrils which often coincide with its core [23, 8, 24]. These spectra allowed us to identify the amino acid residue composition of the core, which included Ala, Asn, Gln, His, Ile, Ser, Pro, Thr, and Val. To evaluate how the structure of these fibrils compared to those extracted from *Drosophila* by Hervás and co-workers, we predicted the NMR chemical shifts of Orb2B residues 178-204 of their cryo-EM structure (PDB access code 6VPS) using the program Shiftx2 [25] and projected the expected cross peaks onto our spectra. As can be seen from Figure 7, the predicted Gln Cα-Cβ, Cα-Cγ, and His Cα-Cβ peaks overlap well with the broad peak at about 55 and 32 ppm. In addition, the predicted cross peak of L198, in the center of the Orb2 core, overlapped perfectly with the Ile Cα-Cβ and Cα-Cγ signals identified in our spectrum. Overall, these data suggest that the Orb2 fibril core determined by cryo-EM is also present in our fibrils. In addition, we identified many intense signals from residues that are not part of the Gln/His rich core namely Ala, Asn, Ile, Pro, Thr, and Val. The chemical shift of these resonances is generally compatible with an extended β-sheet conformation.

**Figure 7:**
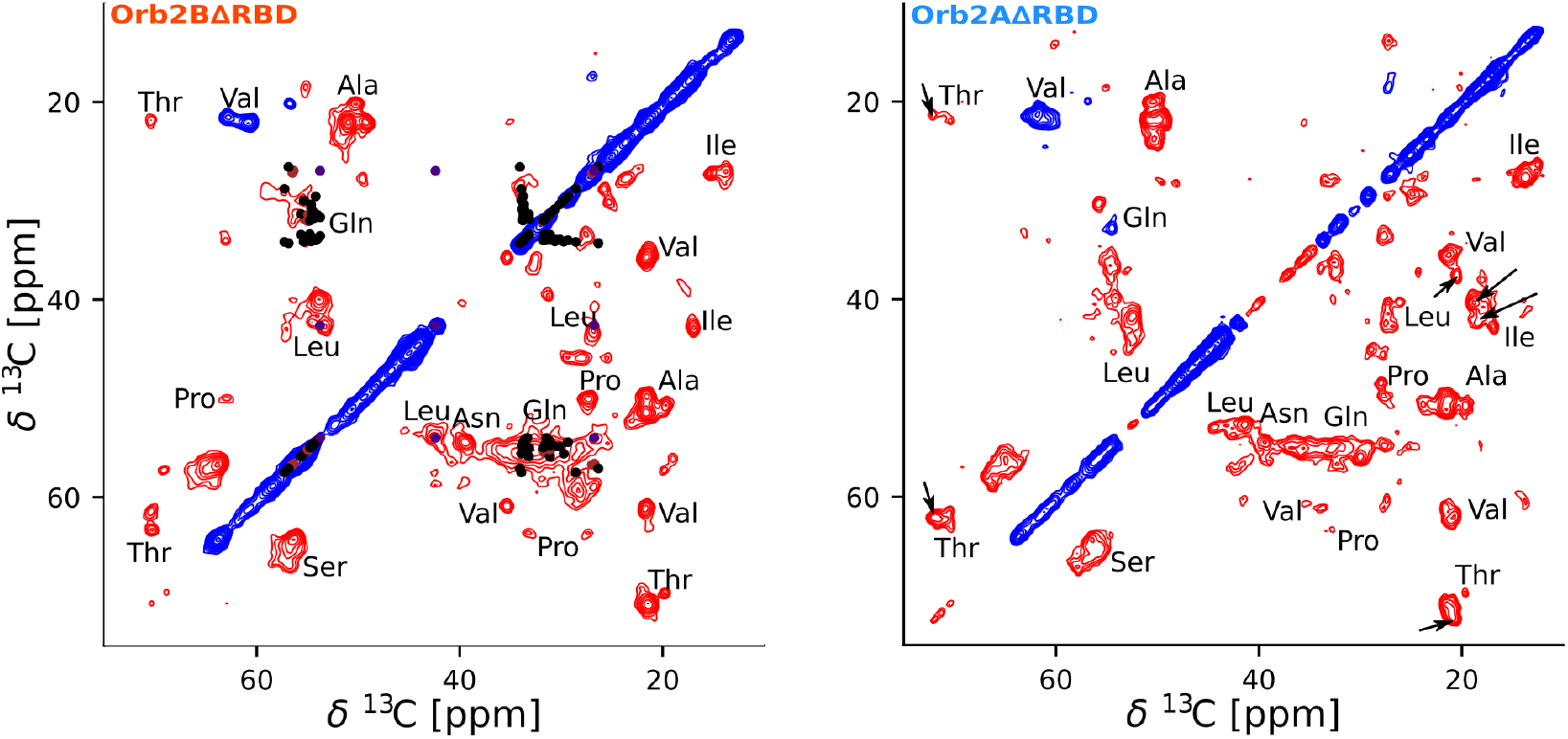
Both Orb2BΔRBD and Orb2AΔRBD fibril spectra show additional immobilized regions beside the Q/H rich fibril core. Solid-state NMR 2D ^13^C-^13^C DREAM spectra of Orb2BΔRBD and Orb2AΔRBD fibrils. Positive signals are shown in blue, negative signals in red. Amino acids type assignments are indicated. Chemical shifts predicted from the cryo-EM structure (6VPS) using the program Shiftx2 are shown in black, brown, and indigo for Gln, His, and Leu, respectively. The good overlap of these predicted chemical shifts with the experimental data indicates the presence of Q/H-rich Orb2 fibril core. The additional intense and narrow signals of amino acids that are not found in the core (e.g. Val, Pro, Ala, Thr, Asn, Ile), suggest that there are static, well ordered regions other than the Q/H rich core in these fibrils. Both spectra are very similar. The few additional cross-peaks detected in the spectra of Orb2AΔRBD fibrils are indicated with arrows.

The 2D DREAM spectra confirm that the two fibrils give spectra that are very similar, indicating a similar core structure. Nevertheless, we were able to identify few additional cross peaks in the DREAM spectrum of Orb2AΔRBD that could not be seen in Orb2BΔRBD. These resonances are highlighted with arrows in Figure 7 and come from Thr, Val, and Ile residues. Interestingly, these resonances do not clearly point to an additional fibril core located at the N-terminal region unique to Orb2AΔRBD as could be expected, because Thr is not found in this N-terminus. We are currently working on the site-specific assignment of these samples to determine the location and structure of the additional static domains and their differences.

How do the Gln resonances in our Orb2 fibrils spectra compare to Gln resonances found in spectra of HTT_ex1_ and Sup35p, two proteins that form fibrils with a Gln-rich core? Figure S5A shows the overlay of 2D ^13^C-^13^C spectra of Orb2BΔRBD and HTT_ex1_ (Q25) fibrils, both of which were formed through phase separation. This overlay shows that the Gln peaks in Orb2BΔRBD do not correspond well to any of the two major Gln peaks (i.e. Gln A and Gln B) in the HTT_ex1_ fibril spectra [26, 27, 28, 29, 30, 31]. Orb2BΔRBD’s Cα and Cβ chemical shift is compatible with an extended conformation and in between the shifts observed for Gln A and Gln B in HTT_ex1_. Its side chain Cδ shift is significantly higher than what has been observed for Gln A and GlnB in HTT_ex1_ and compatible with the minor Gln population in HTT_ex1_ termed Gln C. This analysis suggests that the Gln conformations in the cross-β cores of Orb2 and HTT_ex1_ are different. We also compared the Gln resonance assigned in the Q-rich fibril core of Sup35p [32] with our spectra (Figure S5B). Many of the Gln Cα-Cβ and Cα-Cγ assignments fit well into the broad Gln resonance observed for Orb2BΔRBD suggesting that they have a similar structure.

## 3. Discussion

In the present study, we show that Orb2 can phase separate into droplets that quickly mature and transition into cross-β fibrils in the absence of the C-terminal RNA binding domain. Because Orb2 has a domain structure similar to other RNA binding proteins that phase separate, droplet formation of Orb2 was predicted to occur [10]. However, Majumdar and co-workers showed that intracellular Orb2A puncta showed little mobility when assessed using FRAP assays suggesting that these were self-assembled oligomers rather than droplets [4]. We reconcile these two views by showing that Orb2 phase separation does occur. However, in contrast to other proteins that undergo slow maturation from more liquid to more glass-like droplets with little to no protein mobility, the Orb2 droplets we describe either never go through a liquid phase or this state is so short-lived that we were not able to detect it. In this sense Orb2 droplets are similar to those formed by the nuclear pore complex protein nucleoporin [33], the protein velo1 found in Balbiani bodies [34], or stress induced A-bodies [35, 36]. In contrast, HTT_ex1_ (Q25), which can form protein droplets that mature into fibrils with Gln-rich core similar to Orb2, forms initially liquid-like assemblies [16].

In our hands, phase separation is a necessary requirement to go from soluble Orb2ΔRBD into amyloid fibrils. We previously showed that a shorter fragment of Orb2A (i.e. Orb2A_1−88_) forms cross-β fibrils over time in HSU-buffer [8]. However, the fibril core of Orb2A_1−88_ was not located in the Q/H-rich domain but rather at the very N-terminus unique to Orb2A. In contrast, Orb2ΔRBD fragments did not aggregate in HSU-buffer and required phase separation for fibril formation. Further, NMR spectra of Orb2ΔRBD fibrils are compatible with the presence of the Q/H-rich fibril core described in the recent cryo-EM structure of Orb2 fibrils [6]. Our NMR spectra also show that there are additional static domains in Orb2BΔRBD and Orb2AΔRBD fibrils not explained by the Q/H-rich fibril core. These domains contain Ile, Pro, Val and multiple Ala, Ser, and Thr residues. This amino acid composition does not allow us to locate these domains within the sequence. There is little evidence that they are located in the first 8 residues of Orb2A because the prominent Phe peak that is characteristic for fibrils formed by these residues [8] was missing from our spectra. We are currently working on the assignment of these additional regions.

In addition to having slightly different fibril cores, Orb2BΔRBD and Orb2AΔRBD also form droplets that vary in size independent of conditions. Orb2AΔRBD fibrils are highly bundled, whereas Orb2BΔRBD fibrils show little tendency to bundle. Consequently, the different Orb2 N-termini must play a role in droplet size and fibril bundling.

How do the NMR spectra of Orb2ΔRBD fibrils compare to the NMR spectra of HTT_ex1_ and Sup35p fibrils? This comparison is of interest because the Q/H-rich fibril core of Orb2 is reminiscent of the polyQ fibril core of HTT_ex1_ and the Gln-rich core of Sup35p. Our comparison of the Gln cross peaks of HTT_ex1_ and Orb2BΔRBD fibril spectra indicates that the conformation of the Gln residues in both fibril cores is distinct, which would be in line with the fact that the Orb2 fibril core is an in-register parallel β-structure [6] whereas HTT_ex1_ is not [37]. This might also explain why the assignment of Gln resonances of Sup35p fibrils [32] fits our spectra better than HTT_ex1_, since Sup35 fibrils were shown to form in-register parallel β-sheets [38]. Ultimately, the best fit to the Gln resonances in our spectra was obtained by predicting chemical shifts from the cryo-EM structure of the Orb2 fibril core, suggesting that our fibrils are similar in structure to those found in vivo.

Where Orb2 fragments without the C-terminal RBD form cross-β fibrils after droplet formation, full-length Orb2B remains in a state of matured droplets, which merge but neither completely fuse nor transition into cross-β fibrils. This observation is compatible with previous in vivo work showing that Orb2B alone cannot form functional aggregates that activate the translation of target mRNA [4]. Because Orb2BΔRBD forms fibrils via droplet formation, the C-terminal RBD likely plays an important role in preventing fibril formation of Orb2B.

In summary, our data establish droplet formation as a possible mechanism for inducing the Orb2 fibril formation in vivo. However, this mechanism needs to be confirmed in vivo. Based on our ability to make monomeric Orb2 and induce its phase transition, our future work aims at building a functional Orb2 protein complex in vitro and describe the structural interactions of its components.

## 4. Methods and Materials

### 4.1. Protein constructs

The sequences for Orb2AΔRBD and Orb2BΔRBD were codon optimized for expression in *E. coli*, synthesized, and cloned into pET28b expression vectors with a C-terminal 6x histidine-tag (6xHis-tag) by Genscript USA Inc. Full-length Orb2B was cloned from a pDEST vector (provided by the lab of Dr. Kausik Si) into a pET28b vector.

### 4.2. Protein expressions and purification

#### 4.2.1. Orb2

The Orb2B was expressed in Rosetta 2 *E. Coli* (DE3) whereas, Orb2AΔRBD and Orb2BΔRBD were expressed in T7 express or BL21 (DE3) (EMD Millipore, Billerica, MA). A single colony was used to inoculate 50 ml LB medium with the appropriate antibiotic (chloramphenicol 35 mg/ml and kanamycin 50 mg/ml) and grown overnight at 30°C. The next day, 1-5 ml of this culture was expanded into 500 ml LB with the appropriate antibiotic and cells were grown at 37°C to an OD_600_ of 0.6. Protein expression was subsequently induced with the addition of 1 mM IPTG and cells were grown for an additional 18-20 h at 18°C. Cells were then harvested via centrifugation in a Sorvall SLC-6000 rotor (Thermo Fisher Scientific) at 4000 rpm for 20 min at 4°C and were used immediately or stored at −80°C for future use.

Cell pellets ranging from 2.5-3 g were resuspended in HSU buffer (1 M urea, 100 mM KCl, 10 mM HEPES, 0.1% v/v Tween-20 and 0.05% v/v β-mercaptoethanol, pH 7.6) containing 1 mg/ml lysozyme and 2 μl/ml of 10X DNase I. Cells were then sonicated using a Q125 ultrasonic homogenizer (QSonica, Newton, CT) and the cell lysate was centrifuged at 20,000 rpm for 20 min at 4°C using a Sorvall SS-34 rotor (Thermo Fisher Scientific). The supernatant was collected and filtered through a 0.22 μm vacuum filtration system (Corning INC). The 6xHis-tagged protein samples were loaded onto pre-equilibrated Ni^2+^ affinity resin (HisTrap HP 5 ml, GE Healthcare). The column was sequentially washed with HSU buffer containing 0.5% triton-X, 500 mM NaCl, 20 mM imidazole, and 1 M NaCl (each wash was 5 column volumes). The protein was then eluted with 500 mM imidazole and 1 M NaCl in HSU buffer. Protein elution was traced using 280 nm UV absorption. Fractions containing the protein of interest were pooled together and loaded onto a pre-equilibrated size exclusion column (HiPrep 16/60 Sephacryl S-300, GE Healthcare). The protein was then eluted using HSU buffer and the protein fractions were used immediately or frozen in liquid nitrogen and stored at −80°C.

#### 4.2.2. HTTex1

HTT_ex1_ (Q25) was expressed and purified as described earlier [27, 39]. Reversed-phase purified and lyophilized HTT_ex1_ (Q25) was dissolved in 0.5% trifluoroacetic acid in methanol (TFA-MeOH) and dried in a borosilicate glass tube using nitrogen. The dried protein film was reconstituted in 20 mM phosphate buffer, pH 7.4, containing 150 mM NaCl and 10% dextran such that the final protein concentration was 300 μM. The solution was incubated at room temperature for 14 hours. Under these conditions, HTT_ex1_ (Q25) forms fibrils via phase separation. These fibrils were harvested by centrifugation at 40,000 rpm for 30 min in a Beckman Coulter TLA-100.3 rotor. The fibrils were washed with 20 mM phosphate buffer, pH 7.4, containing 20 mM NaCl before they were packed into a solid-state NMR rotor.

### 4.3. Fluorescence and DIC wide field microscopy

Protein samples that were used for microscopy either eluted from SEC in a HSU buffer that contained 5 mM TCEP or exchanged into a HSU-buffer with 5 mM TCEP using a desalting column. The protein samples were then labeled with either 5 mM thiol-reactive or 5 mM amine-reactive fluorescent dyes over night at 4°C with light shaking. Orb2B and Orb2BΔRBD were labeled with thiol-reactive Oregon Green 488 through maleimide chemistry (Thermofisher). Due to the lack of cystine residues in the Orb2AΔRBD protein construct, it was labeled with lysine-reactive succinimidyl ester Oregon Green 488 (Thermofisher). The excitation and emission maxima of these fluorescent bioconjugates are 496 nm and 524 nm, respectively. To induce droplet formation, protein was either exchanged into H-buffer (10 mM HEPES, 0.03% v/v NaN_3_ and 0.05% v/v β-mercaptoethanol, pH 6.5), or into HSU buffer, pH 6.5 followed by the addition of 10% v/v PEG 8000 or 1:10 m/m PolyA:protein.

Fluorescence images were taking by exiting at 488 nm and imaging at 505 nm using a Zeiss AxioImager widefield light microscope with white light laser. DIC microscopy images were also recorded on Zeiss AxioImager microscope seconds after the corresponding fluorescence images were taken, by switching from a Zeiss Filter set 38 HE (optimized for emission at 488 nm) to a default Zeiss DIC filter. Image of protein droplets were captured at 0, 12 and 48 hours post exchanged into the H-buffer. Images were analyzed using Fiji image analysis software [40, 41] including the BioVoxxel plugin. Microscopy was repeated on three biological replicates.

### 4.4. Fluorescence recovery after photobleaching (FRAP)

Oregon Green labeled protein samples were prepared analogous to those described above. Protein samples were then exchanged into H-buffer with 5 mM TCEP using a PD-10 G-25 media desalting column and FRAP measurements were done using Leica-SP8X confocal laser scanning microscope with a 63x oil immersion objective. Droplets were bleached with an average diameter of 5 nm, to avoid whole-droplet bleaching artifacts in droplets of smaller diameters [42]. Bleached regions of interest (ROI) of 0.5 μm were created with a pinhole Argon laser with a scanning speed ranging from 50-150 ms using the Leica software. FRAP data were analyzed and plotted with in house python scripts using the pandas, numpy, scipy, and matplotlib libraries. Recovery curves *±* SD were generated by normalizing and averaging across three biological replicates.

### 4.5. Thioflavin T fluorescence assay

Protein samples were exchanged into a H-buffer using a PD-10 G-25 media desalting column (GE Healthcare) to induce droplet formation. 200 μl of 10-20 μM protein were mixed with thioflavin T (Tht) at a final concentration of 50 μM and added to a clear flat bottom 96-well plate (Greiner Bio-One). The solution was then excited at 442 nm, and the emission at 482 nm was measured every 10 minutes for 80-160 hours using an Eppendorf AF2200 plate reader at 25°C. Three independent measurements were recorded for each protein, and normalized and plotted using in house python scripts (available upon request).

### 4.6. Electron Microscopy

Electron microscopy specimens were prepared by submerging a Formvar carbon coated copper grid (Electron Microscopy Sciences) in 10 μl of protein solution and incubating it for 5 minutes. These grids were then stained with 10 μl of a 1% uranyl acetate solution for 2 minutes at room temperature. The grids were then tapped on two additional 10 μ1 uranyl acetate drops and finally, rinsed in deionized water. Filter paper was used to remove the excess of stain and water before imaging. The negatively stained samples were examined for droplets or fibrils with a JEOL JEM-1400 TEM electron microscope. Images were acquired using a Gatan Orius digital camera at magnification of 5000-10,000X. Fibril images were analyzed using ImageJ including the FibrilJ plugin [43].

### 4.7. Solid-state NMR spectroscopy

NMR spectra were recorded on an Agilent DD2 600 MHz solid-state NMR spectrometer using a T3 1.6 mm probe. All hard ^1^H and ^13^C radio frequency (rf) pulses had amplitudes of 200 kHz and 100 kHz, respectively. ^1^H-^13^C cross polarizations (CPs) were done with 60 kHz rf-amplitude on ^13^C and a ^1^H rf-amplitude that was larger by the MAS frequency. A 10% amplitude ramp was applied during 1 ms of contact time. Proton decoupling during acquisition was done using the XiX decoupling scheme [44] with rf-field amplitudes of 140 kHz. A recycle delay of 3 s and a 0°C set temperature was used for all spectra.

One dimensional CP, DB, and refocused INEPT spectra were recorded at 25 kHz MAS with a spectral width of 50 kHz, 1000 complex points, and 1024 acquisitions. ^13^C-^13^C DREAM (dipolar recoupling enhanced by amplitude modulation) [45] spectra were recorded at 30 kHz MAS using a spectral width of 50 kHz in both dimensions and a mixing time of 4.5 ms. For each of the 800 indirect TPPI increments 128 and 64 acquisitions were recorded for the Orb2BΔRBD and Orb2AΔRBD samples, respectively.

2D DARR (dipolar assisted rotational resonance) spectra [46] of Orb2BΔRBD and HTT_ex1_ (Q25) were recorded at 25 kHz MAS with a spectral width of 50 kHz, and a mixing time of 50 ms. For Orb2BΔRBD 400 indirect, complex increments were recorded with 64 acquisitions each. For HTT_ex1_ (Q25) 500 indirect, complex increments were recorded with 64 acquisitions each.

All spectra were processed using Lorentz to Gauss transform window functions. Adamantane spectra were used to reference the chemical shifts externally to DSS (4,4-dimethyl-4-silapentane-1-sulfonic acid) [47]. The spectra were analyzed using CARA [25] and plotted using in house python scripts (available upon request) based on the numpy, matplotlib, and nmrglue packages [48, 49].

## 5. Acknowledgments

A.B.S. would like to acknowledge funding from the National Institutes of Health (R01NS084345, R01GM110521) and the CHDI Foundation (Award A-12640). A.S.F. would like to acknowledge funding from the National Institutes of Health (F31GM120858).

## 8. Supporting Figures

**Figure S1:**
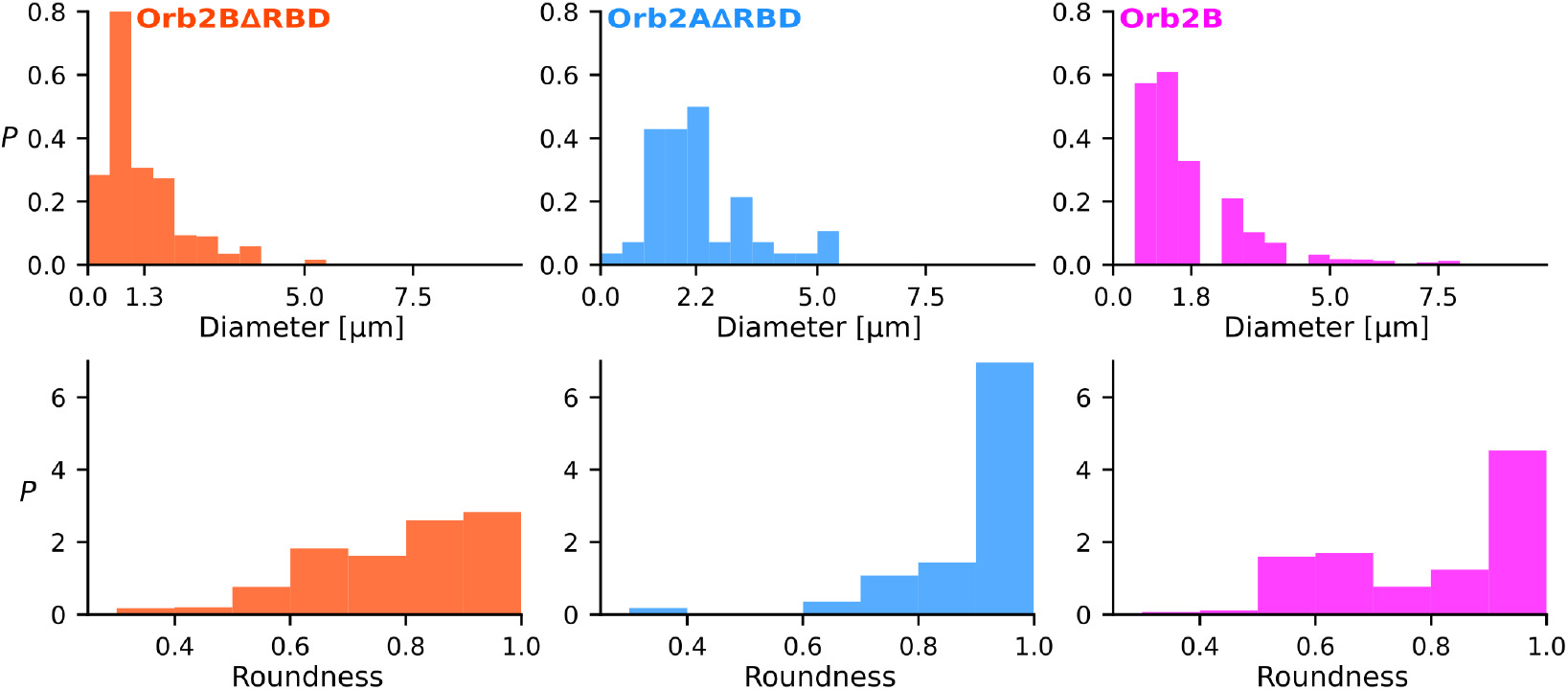
Orb2 droplets are generally between 0 and 5 μm in diameter and have a high degree of roundness. Diameter and roundness of Orb2 droplets were determined using the ImageJ BioVoxxel plugin. Histograms are presented as probability density *P* and the average diameter is indicated.

**Figure S2:**
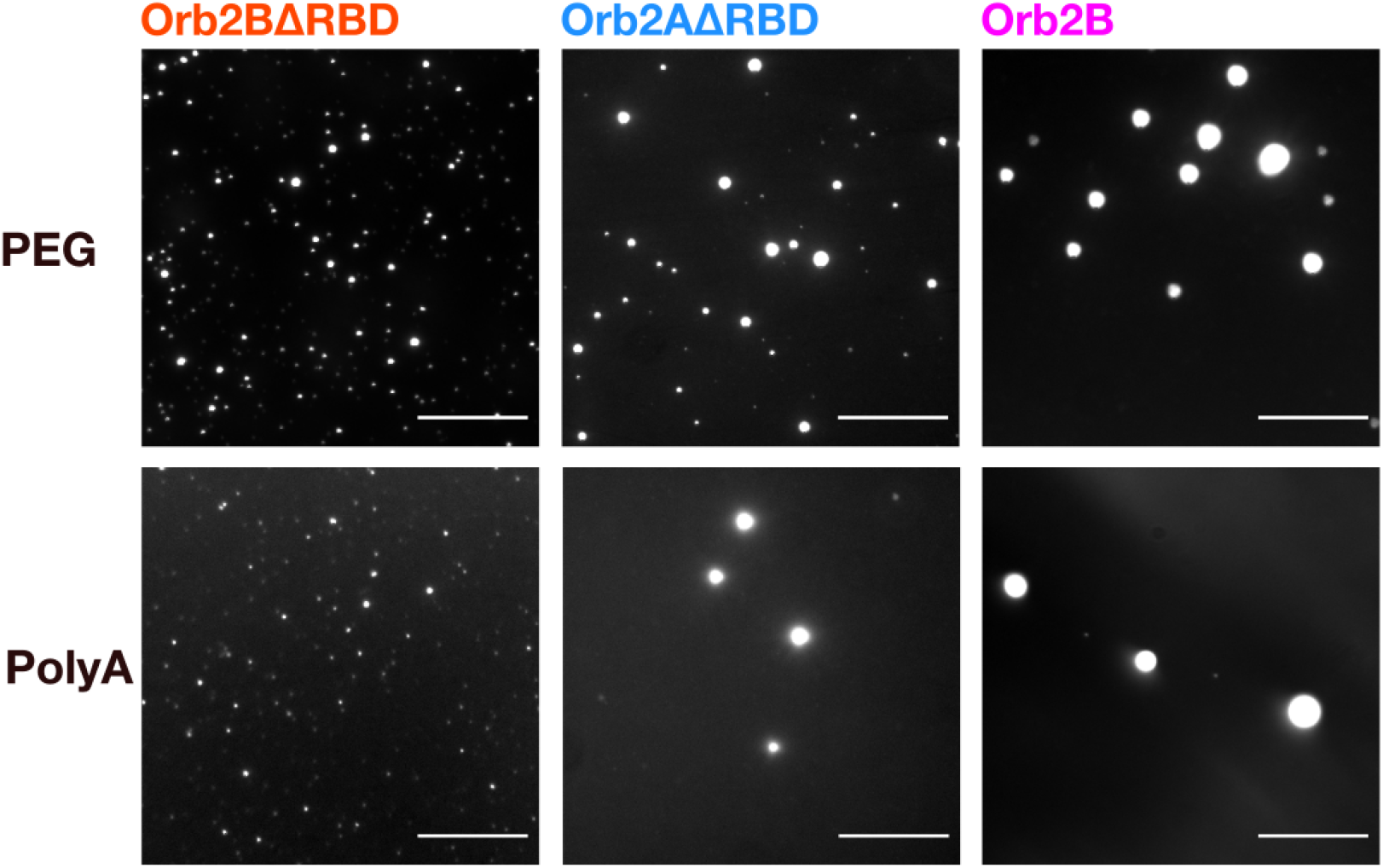
Orb2 also forms droplets in the presence of PEG and RNA. Fluorescence microscopy images of droplets formed by Orb2BΔRBD, Orb2AΔRBD, and Orb2B immediately after adding 10% v/v PEG 8000 or PolyA at a 1:10 PolyA:protein m/m ratio. Scale bars represent 20 μm.

**Figure S3:**
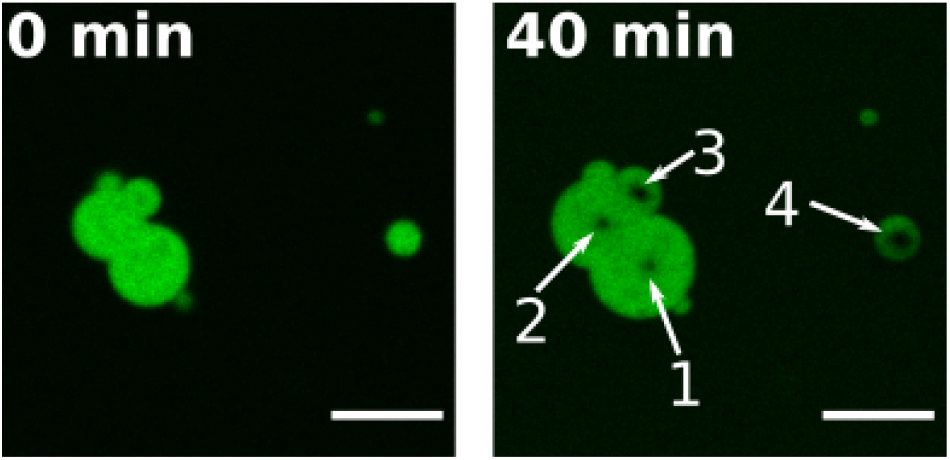
Orb2B shows neither FRAP recovery nor droplet fusion after 40 minutes. Merged Oregon Green 488 labeled Orb2B droplets before (0 min) and after 4 consecutive (indicated with numbers 1-4) 10 minute FRAP experiments (40 min). The fact that neither fluorescence recovery nor droplet fusion events could be observed indicates the relatively static nature of Orb2B inside these droplets. Scale bars represent 5 μm.

**Figure S4:**
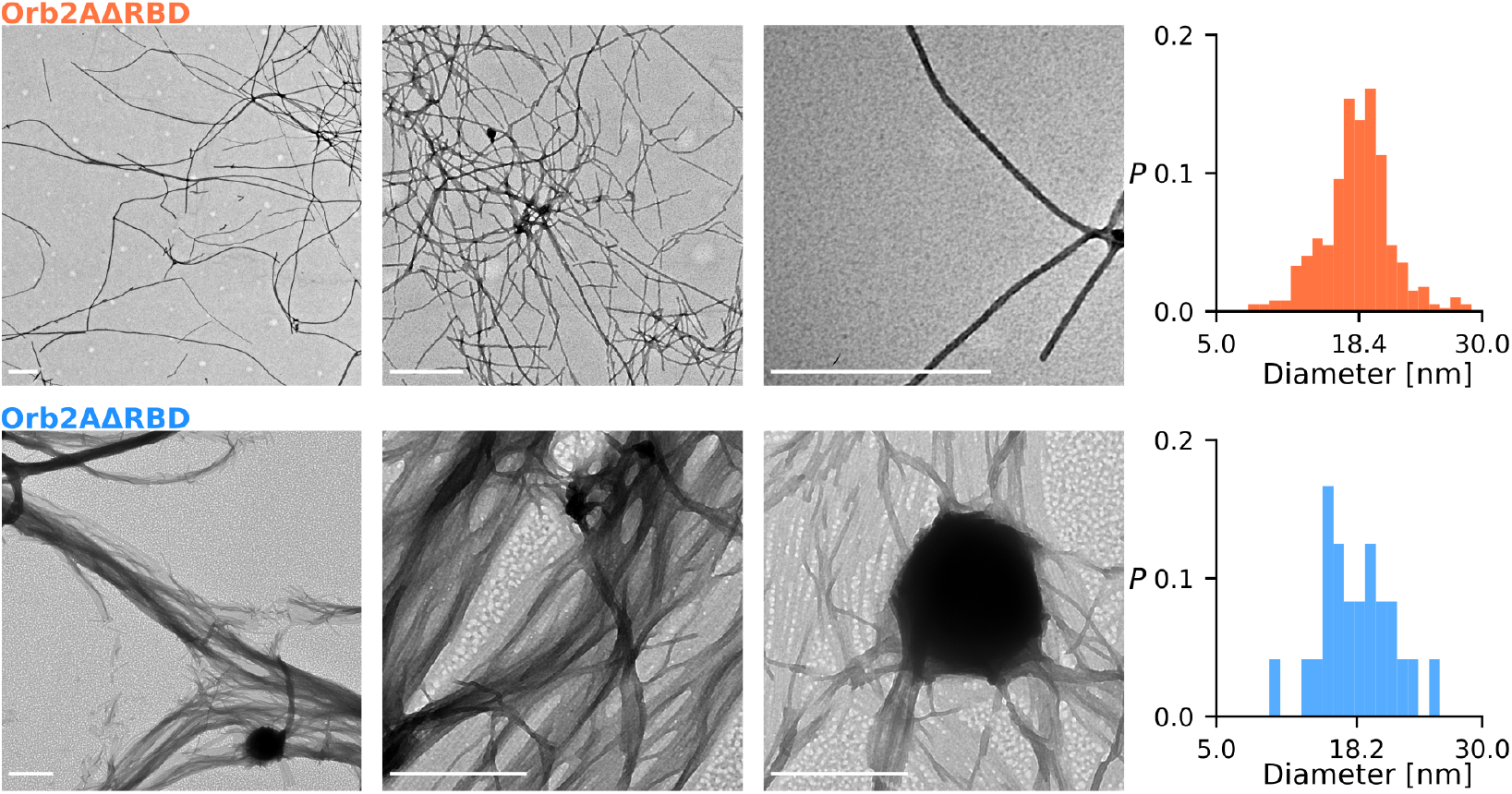
Orb2BΔRBD fibrils are relatively unbundeled, whereas Orb2AΔRBD fibrils are highly bundeled. Both fibrils have a diameter of about 18 nm with no visible twist. Negative stained EM images of Orb2BΔRBD and Orb2AΔRBD fibrils 48 h after buffer exchange. Scale bars denote 500 nm. Fibril diameters were measured using the ImageJ FibrilJ plugin. Histograms are presented as probability density *P* and the average diameter is indicated.

**Figure S5:**
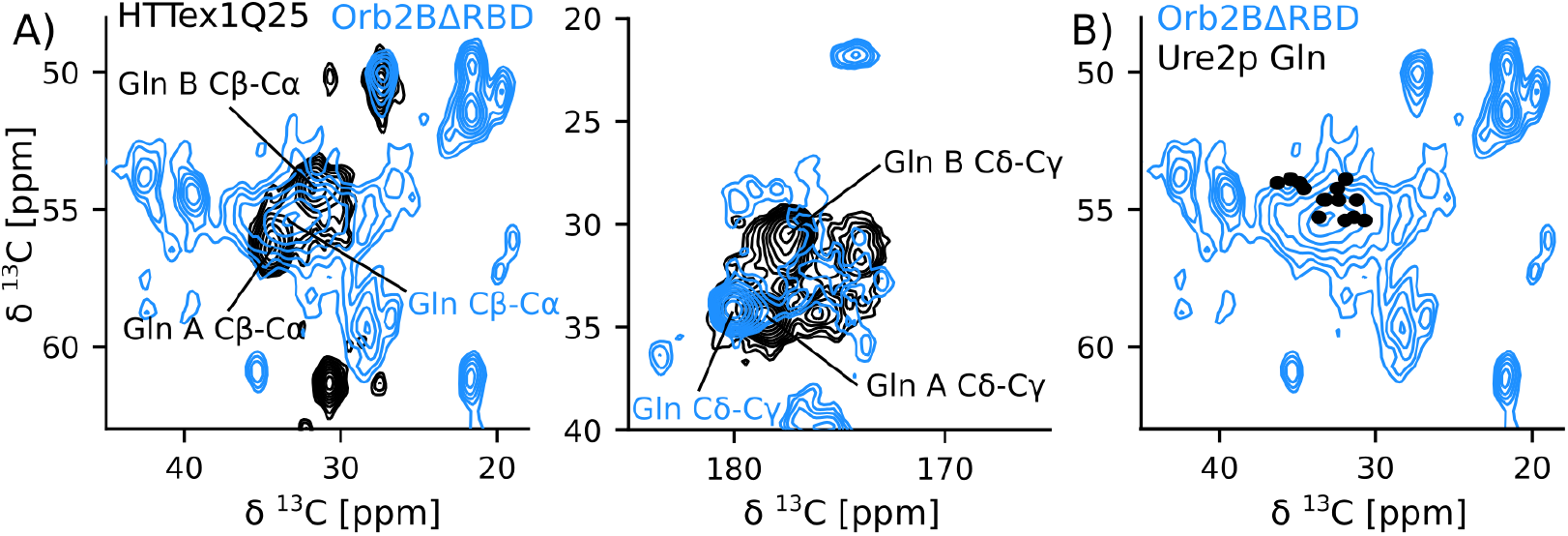
Comparison of Gln peaks in Orb2BΔRBD fibril spectra with HTT_ex1_ (Q25) and Sup35p data. A) Overlay of 2D ^13^C-^13^C spectra of Orb2BΔRBD shown in blue and HTT_ex1_ shown in black. Gln peaks are labeled in the same colors. Where HTT_ex1_ has two major Gln conformations Gln A and Gln B, Orb2BΔRBD has only one. The Gln peak of Orb2BΔRBD does not overlap well with either Gln A and Gln B but its Cδ-Cγ peak overlaps with the minor HTT_ex1_ Gln conformation Gln C. The HTT_ex1_ (Q25) spectrum was recorded using the 2D DARR experiment at 25 kHz MAS with a mixing time of 50 ms. Orb2BΔRBD spectrum shown in left panel is a 2D DREAM recorded at 30 kHz MAS, the right panel shows a 2D DARR recorded at 25 kHz MAS with a mixing time of 50 ms. B) Spectrum of Orb2BΔRBD fibrils with black circles indicating the Gln assignments in the core of Ur2p fibrils (BMRB access code 18407). Although not perfect, the Gln resonances of Sup35p are more similar to Orb2BΔRBD compared to HTT_ex1_.

## Notes

### Competing Interest Statement

The authors have declared no competing interest.

